# Honest and cheating strategies in a simple model of aggressive communication: the role of spatial correlations

**DOI:** 10.1101/257394

**Authors:** Szabolcs Számadó, András Szántó

## Abstract

The evolution and maintenance of communication in terms of aggressive interactions is a long-debated issue. Several game theoretical models and individual based computer simulations investigated this problem in terms of a simple game of aggressive communication. So far all of these investigations focused on well mixed population of individuals. However, spatial correlations can emerge in nature where individuals or group of individuals defend resources. The extensive literature on cooperative games show that these spatial correlations can be vital in the maintenance and evolution of cooperative strategies, thus it is reasonable to expect that such correlations could play an important role in the evolution of honest communication as well. Here we investigate a traditional game of aggressive communication in a spatially explicit context. We investigate the role of spatial correlations by comparing results of evolvability in well mixed populations with results from spatially explicit populations. Spatial correlations seem to inhibit the evolution of communication in the spatially explicit version of this game. This result is unexpected, and it requires further investigation to understand.

## 1. Introduction

The idea that animals could honestly communicate their intentions or other relevant qualities in context of aggressive interactions fascinated biologist for a long time. On the one hand, there is a clear conflict of interest between participants as the goal is to displace the rival from the contested resource. At first it seems that this makes honest communication highly unlikely. Early game theoretical models seemed to confirm this conclusion. On the other hand, animals often use intricate behaviours and morphological features to influence the rival in a non-physical way. Why would animals waste time and energy on these displays if communication is hopeless in the context of aggressive encounters? It turned out, however, that the situation is not entirely hopeless and there are conditions that allow the honest communication of intentions and relevant qualities in the context of aggressive communication. Enquist [1] was the first to show that variation in behaviour can be a reliable signal of intentions or quality.

Subsequently several variants of this model were used to investigate different problems within aggressive communication: whether weak or strong individuals should use handicaps [2]; the existence and stability of mixed cheating maintained by frequency dependent selection [3]; whether threat displays are handicaps [4]; how proximity risk can stabilize threat displays [5]; or how long-term commitment can stabilize dominance displays [6]. The model also inspired a series of computer simulations [7–9] investigating the evolutionary trajectories and evolutionary stability of honest and cheating equilibria in terms of aggressive communication. Her we investigate a spatial version of the model to study the effect of spatial correlations on the evolution of honest and mixed cheating equilibria.

## 2. The Model

Here we use a slightly modified version of Enquist’s model [1]. Enquist’s model can be seen as a modified version of the Hawk-Dove game [10]. Since the original model was publish with a specific question in mind it is underspecified in other regards. Accordingly, it needs additional assumptions outisde of the original question. subsequent investigations used various modifications, here we use the pay-off and the full strategy set as suggested by Helgesen et al. [11]. The model is a two-person game of aggressive communication where individuals compete for the possession of an indivisible resource. Individuals differ with regard of one relevant quality, which can be called conveniently as “strength”, but it can represent any property relevant in the given context. Each player can be weak or strong and knows its own strength but not that of the opponent. It is a two-stage game where the first stage is communication and the second stage is fight (see Figure 1). In the first step each player can chose between two cost-free signal A or B, then in the second round each animal can give up, attack unconditionally or attack if the opponent does not withdraw. Let *V* denote the value of the contested resource, and *C_ww_* and C_*ss*_ the expected costs of a fight between two weak and two strong individuals respectively. We assume that a strong animal can always beat a weak one with a cost *C_sw_*, and *C_ws_* is the expected cost that a weak animal should suffer on this occasion. The following relation holds between these costs: *C_ws_* > *C_ww_*, *C_ss_* > *C_sw_.* Let us denote the cost of fleeing as *F_f_* and the cost of attacking a fleeing opponent as *F_A_*, and further suppose that there is a cost of waiting if the opponent attacks unconditionally (*F_p_*). It is biologically realistic but not necessary to assume that C_sw_ > *F_a_* and *C_sw_* > *F_p_* (Hurd, 1997), let us assume in line of our original argument that the cost of fleeing is small thus: *C*_ws_ > *Ff.* Individuals of equal strength have an equal chance winning against each other. The frequencies of weak and strong individuals are denoted by *q* and 1-*q* respectively. Then the payoffs for weak and strong contestants can be written as shown in Table 1.

**Figure 1.**
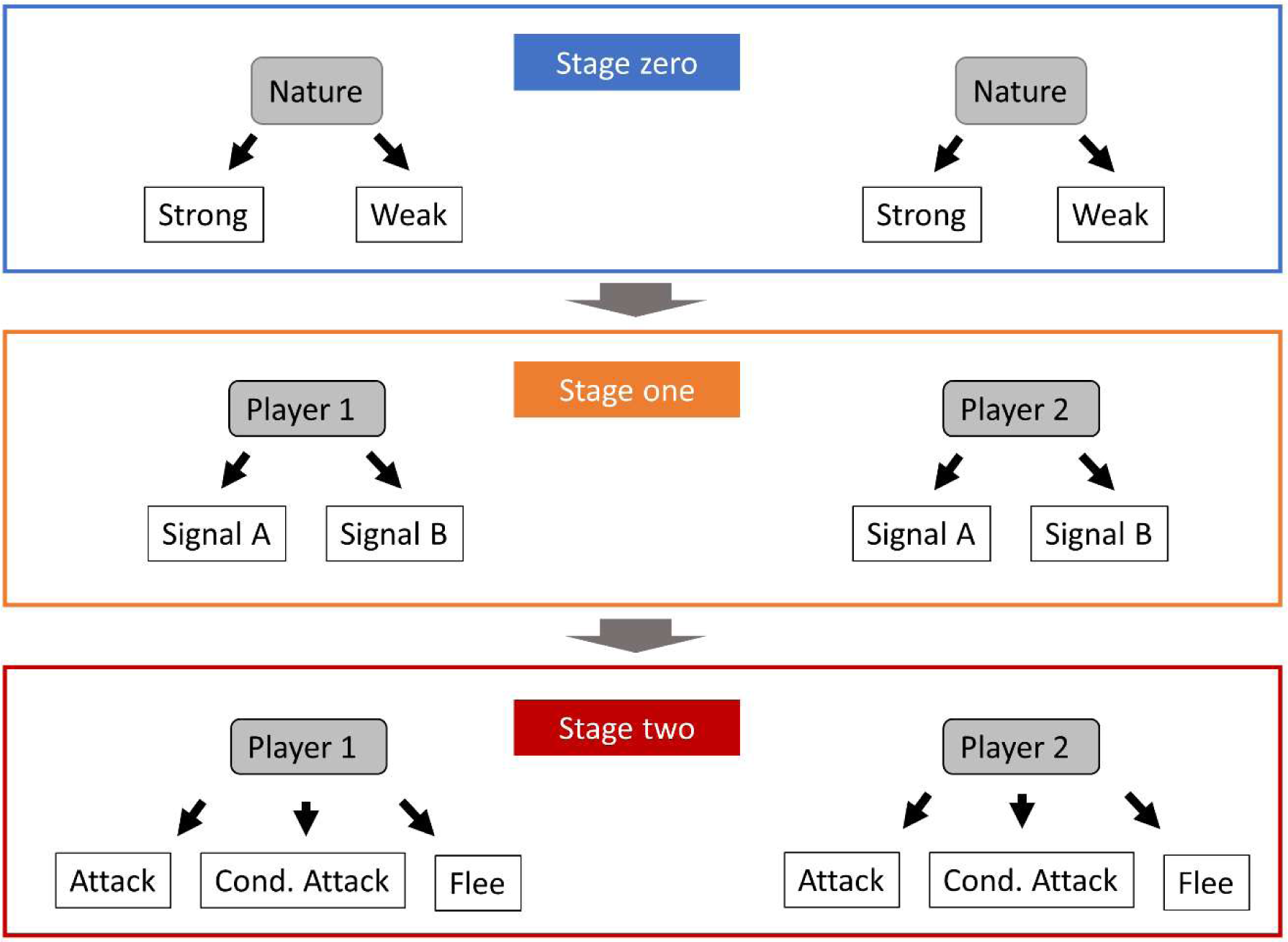
Schematic description of Enquist’s game [1] (after [9]). Stage zero: Nature picks the state of the contestants; this stage is hidden from other players. Stage one: each contestant picks a signal, A or B. Stage two: each contestant picks a behaviour as a response to the signal: Flee, Conditional Attack or Attack.

**Table 1.** Payoffs matrix (after [3]).

| Ego Strength |  | Opponent Strength |  |  |  |  |  |
| --- | --- | --- | --- | --- | --- | --- | --- |
|  |  | Strong |  |  | Weak |  |  |
|  |  | Attack | Cond. attack | Flee | Attack | Cond. attack | Flee |
| Strong | Attack | $0.5V - C_{SS}$ | $0.5V - C_{SS}$ | $V - F_A$ | $V - C_{SW}$ | $V - C_{SW}$ | $V - F_A$ |
| | Cond. attack | $0.5V - C_{SS} - F_P$ | $0.5V - C_{SS}$ | $V$ | $V - C_{SW} - F_P$ | $V - C_{SW}$ | $V$ |
| | Flee | $-C_{SS}$ | 0 | $0.5V$ | $-C_{SW}$ | 0 | $0.5V$ |
| Weak | Attack | $-C_{WS}$ | $-C_{WS}$ | $V - F_A$ | $0.5V - C_{WW}$ | $0.5V - C_{WW}$ | $V - F_A$ |
| | Cond. attack | $-C_{WS} - F_P$ | $-C_{WS}$ | $V$ | $0.5V - C_{WW} - F_P$ | $0.5V - C_{WW}$ | $V$ |
| | Flee | $-C_{WS}$ | 0 | $0.5V$ | $-C_{WW}$ | 0 | $0.5V$ |

Note, while Enquist original model and some of the subsequent investigations used and exogenous explanation for the origin of variation in strength [1, 7, 11], other studies studies investigated populations with endogenous variation, i.e. where variation in strength is maintained by frequency dependent selection [3, 8, 9]. Note that endogenous explanations require some change in the modelling assumptions (see [3]) otherwise the polymorphism of state cannot be maintained. While exo- and endogenous explanations are equally valid, here we investigate a model with endogenous source of variation in order to be able to compare the results with a previous study [9].

Because individuals are weak or strong for life, in the current implementation of the model, therefore half of their genes will be never expressed (see Figure 2). When classifying the behaviour of the individuals, these inactive genes can be safely ignored, thus greatly simplifying the analysis. Notably, these inactive alleles are still present (even if they are not used for classification), and they can be turned on by mutation. Accordingly, I will consider only 36 strategies in the further analysis (see Appendix 1).

**Figure 2.**
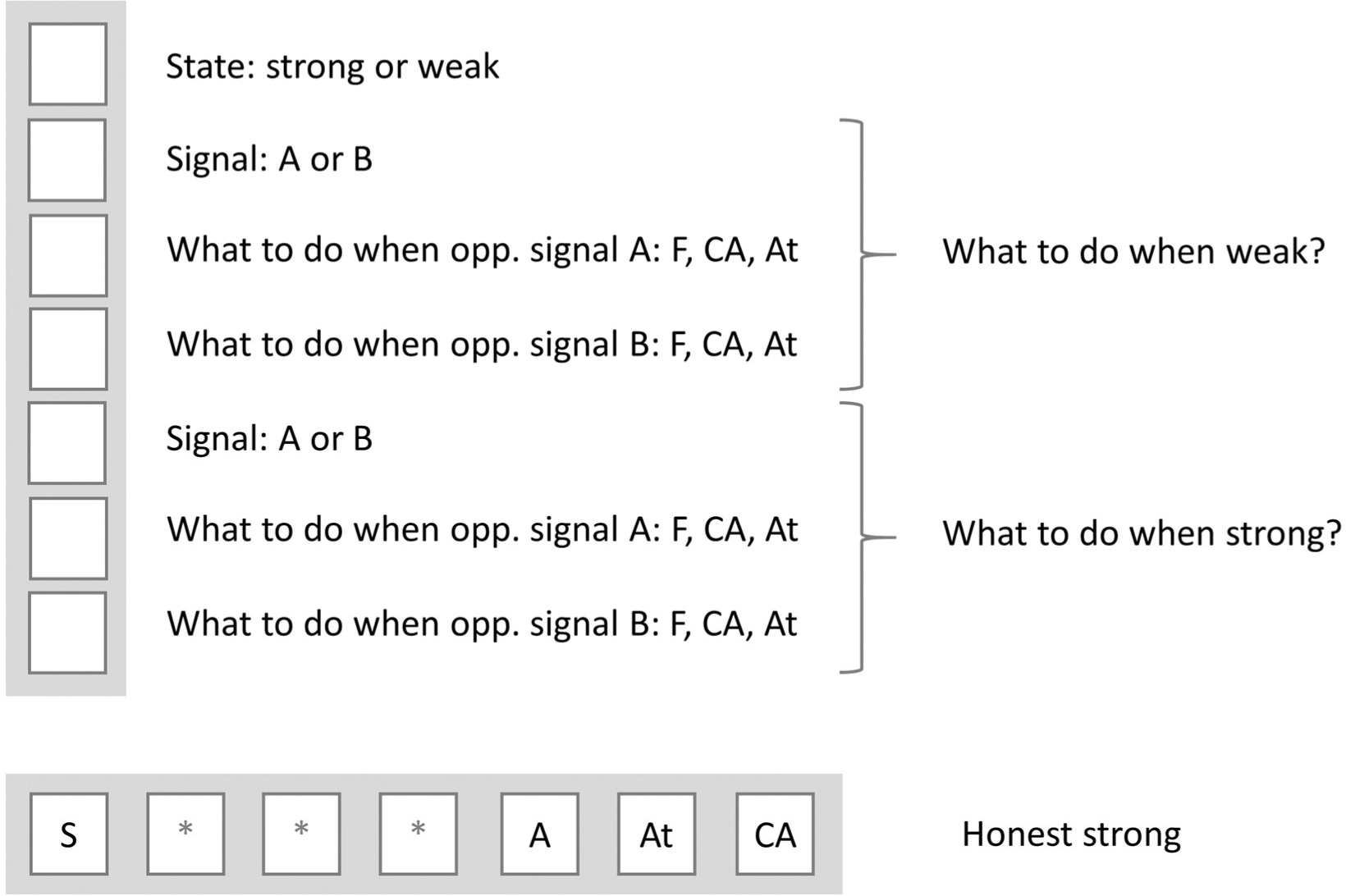
Schematic representation of the coding of the behaviour of individuals (after [9]). Each individual has seven genes: the first gene represents the state of the individual (weak or strong), the next three and the last three encodes the behaviour of the individual depending on whether the state of the individual is weak or strong respectively. Out of these three genes the first gene gives which signal to use; the second gene encodes which behaviour to use as a response to signal A; and finally, the last gene encodes which behaviour to use as a response to signal B. W: weak; S: strong; F: Flee; CA: Conditional Attack; At: Attack. Asterisks denote silent regions which do not influence the behaviour of the given individual. The “genom” of an individual playing the Honest Strong strategy is at the bottom as an example.

Here we use a pairwise comparison rule (sometimes called imitate the better). It is a frequently used update rule in individual based simulations [12–16]. In this update a randomly chosen focal individual’s fitness is compared to one its neighbours. The probability of the neighbour taking over the focal site (*p*) is proportional to the fitness difference between the two individuals if it is positive and zero otherwise. Where f is the maximum potential fitness difference between any two individuals, in order to 0≤p≤1 ([12].

Here we investigate the evolutionarily stability of polymorphic equilibria. We investigate the following types of equilibrium; 50 parameter combinations picked from Szalai and Szâmadô (SS09) [8] in each regions. (i) Honest-strong, Honest-weak (SS09 code: c3; current code, see Appendix 2.: <30,2>); and (iii) Honest-strong, Honest-weak, Liar-weak (SS09 code: c11; current code: <30,2,14>). Supplementary files mixedESS_code3.xls and mixedESS_code11.xls show these combinations.

## 3. Results

No honest communication evolved in the spatially explicit simulations. Mixed cheating evolved in the n=4 scenarios and in the code3 region with n=8. No communication evolved with n=24. The well mixed simulations show a mixture of honest, mixed cheating equilibria as well as no-communication. Figure 3 sums up the results. Changing the neighbourhood size (*n*=4,8,24) had no effect on the results. Figures 4–8 show typical timelines from spatially explicit population. Figures 9–13 shows the final stage of the spatial grids. Videos: grid_387_n8.mp4, grid_8249_n8.mp4, grid_387_n24.mp4, grid_8249_n24.mp4 shows the change of spatial grids.

**Figure 3.**
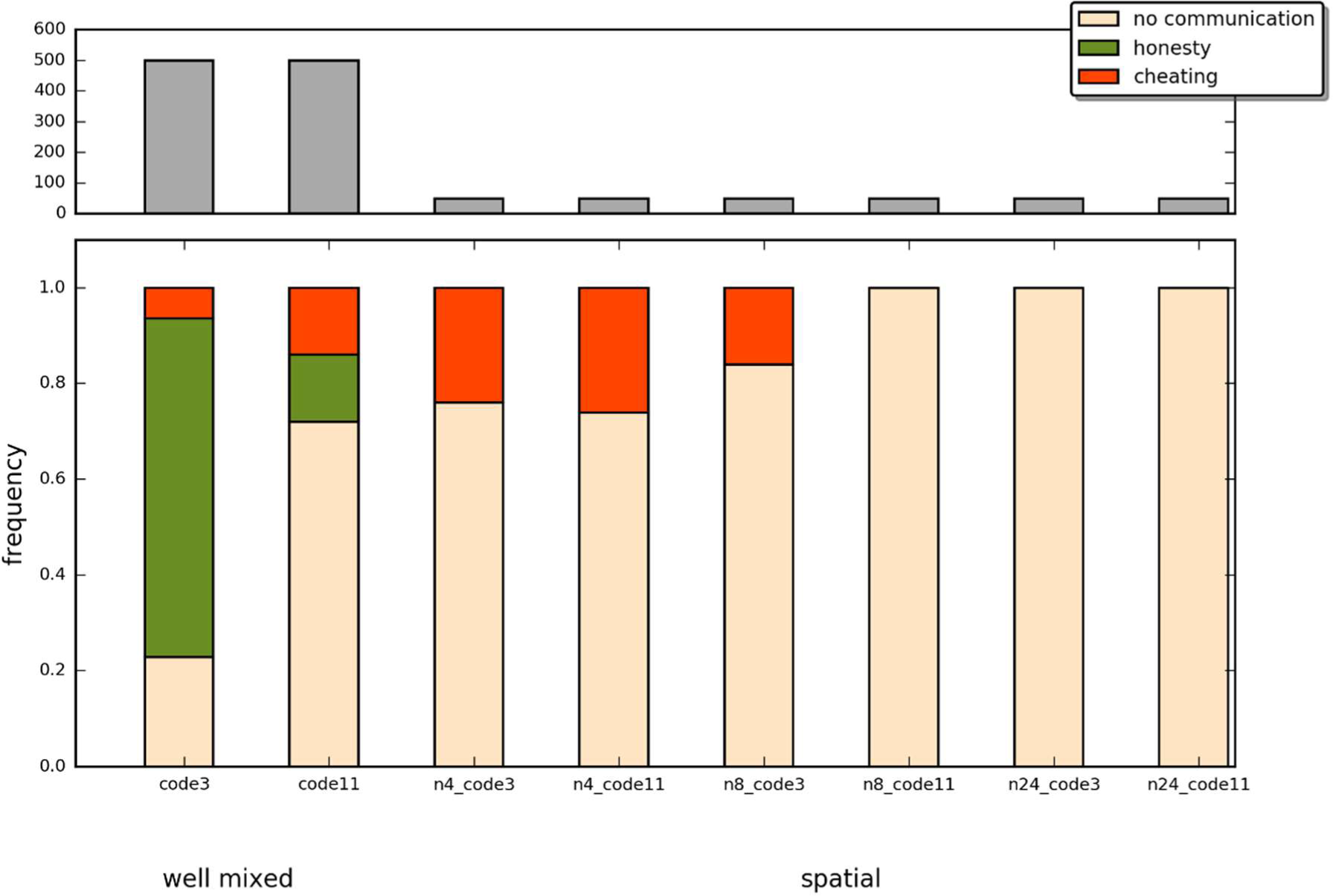
Summary of the results as a function of well mixed vs. spatial runs, neighbourhood size and parameter regions. White: no communication, green: honest signalling, orange: mixed cheating.

**Figure 4.**
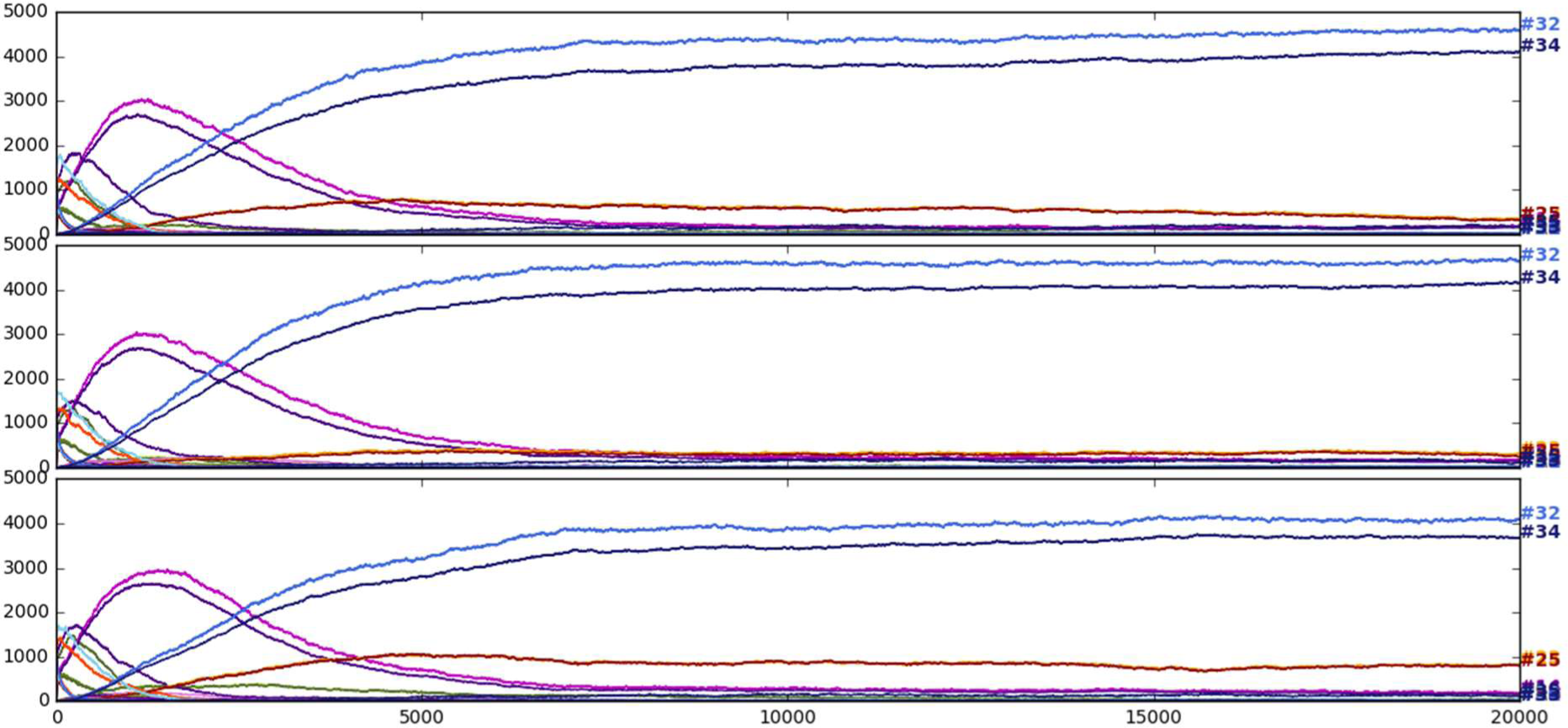
Timeline of three independent runs from region c11, code=8249; neighbourhood size 4.

**Figure 5.**
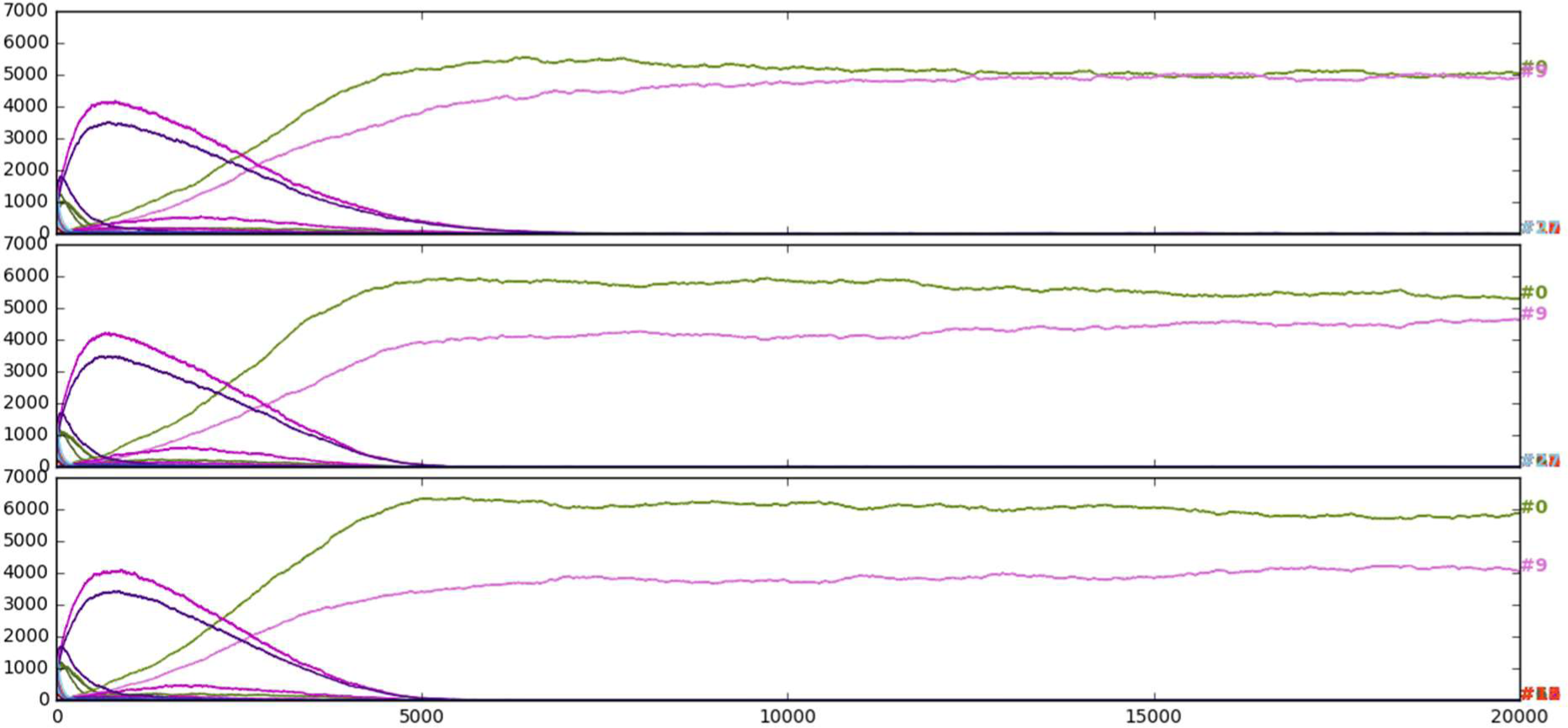
Timeline of three independent runs from region c11, code=387; neighbourhood size 4.

**Figure 6.**
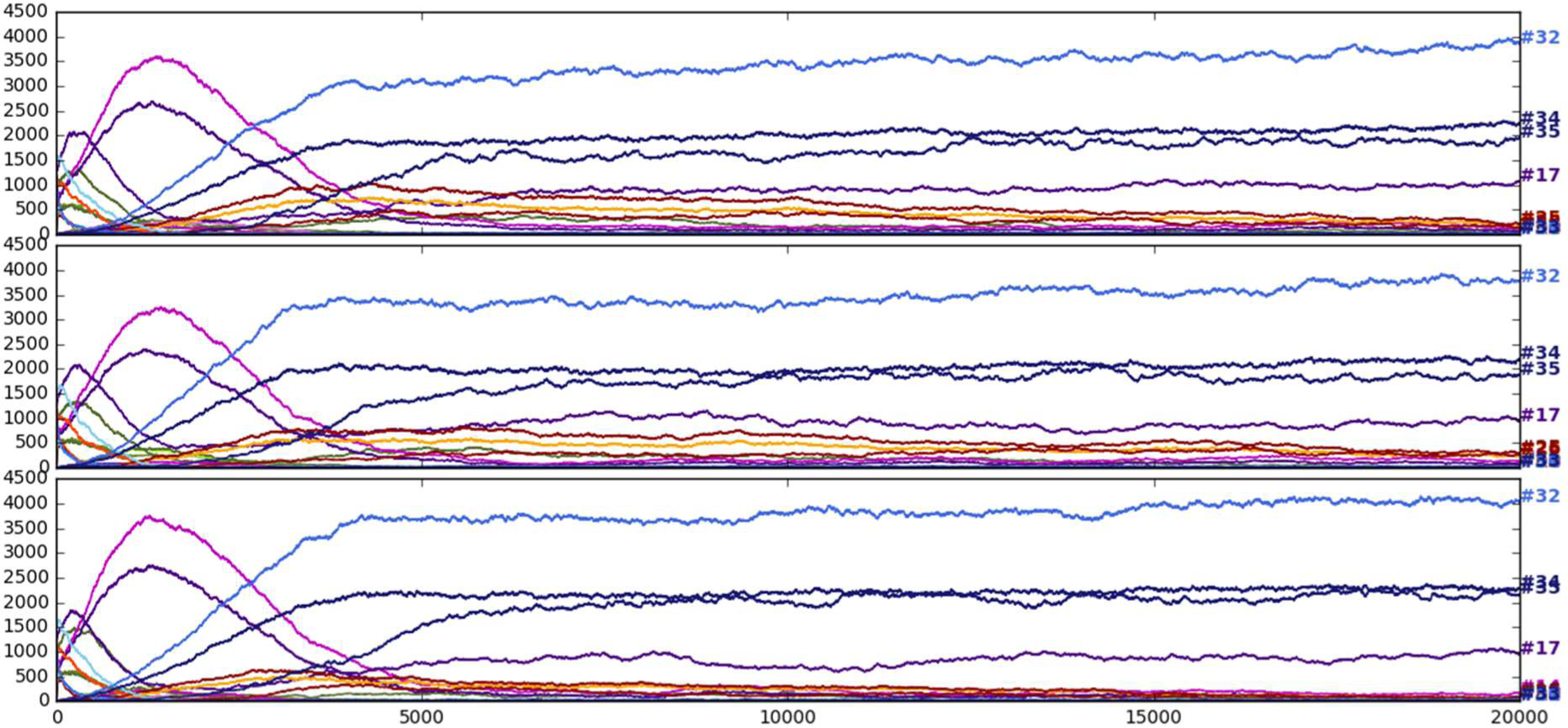
Timeline of three independent runs from region c11, code=8249; neighbourhood size 8.

**Figure 7.**
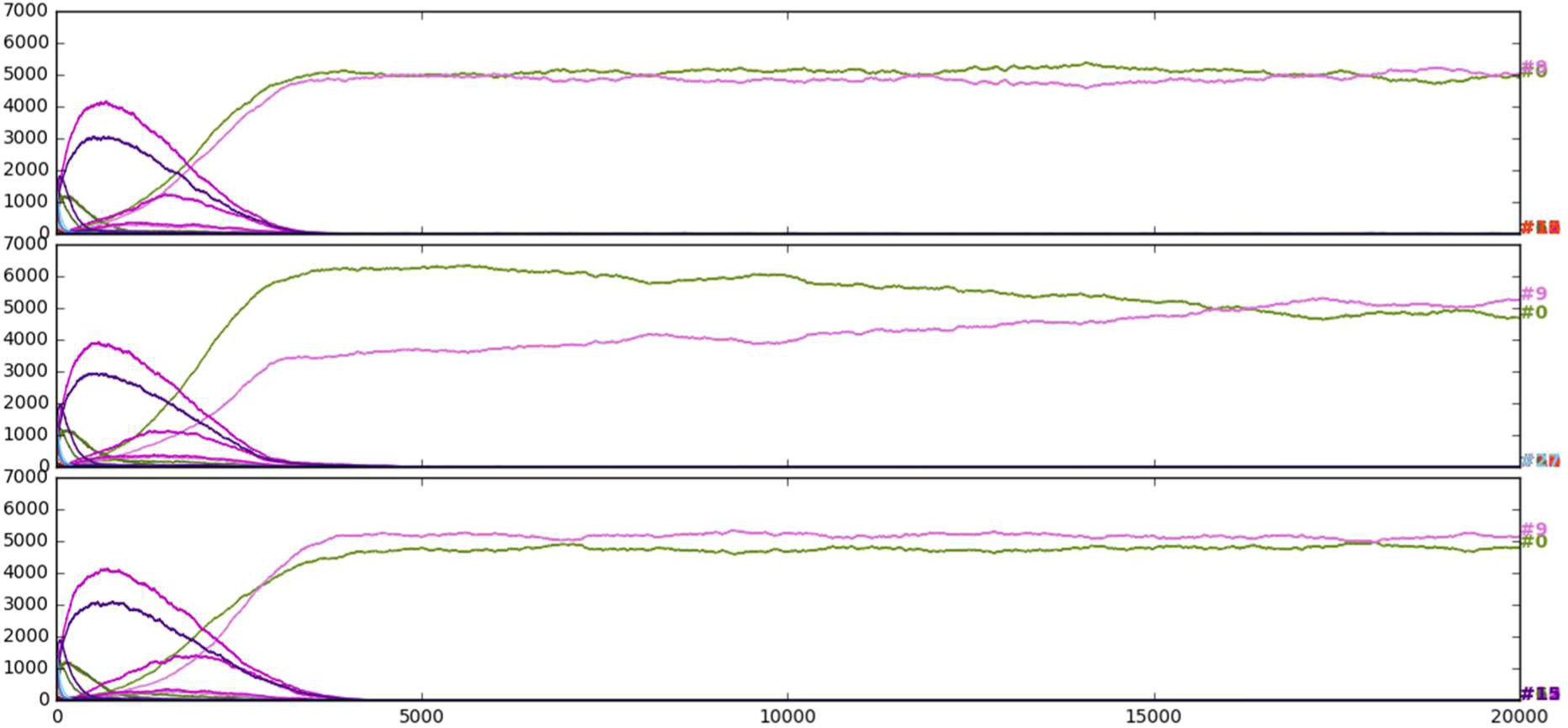
Timeline of three independent runs from region c11, code=387; neighbourhood size 8.

**Figure 8.**
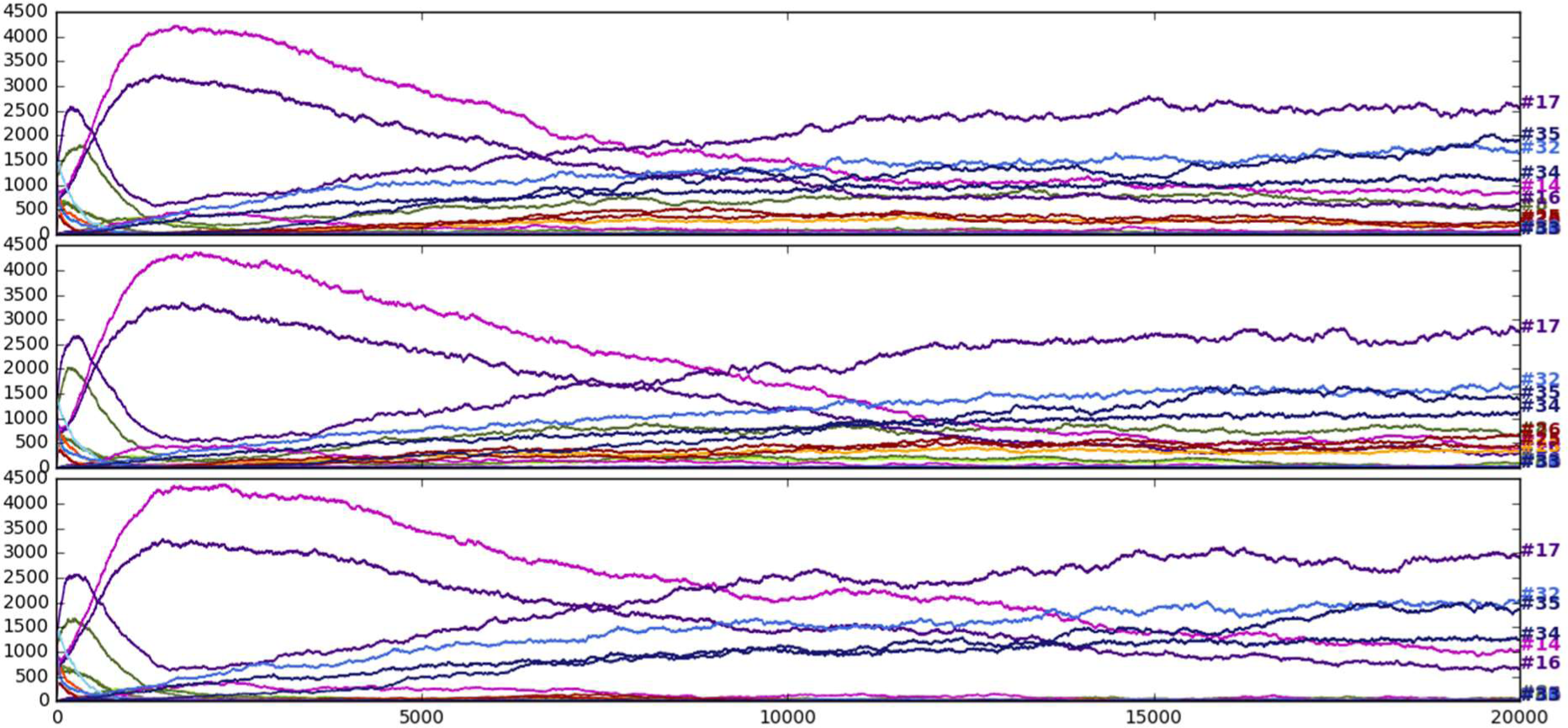
Timeline of three independent runs from region c11, code=8249; neighbourhood size 24.

**Figure 9.**
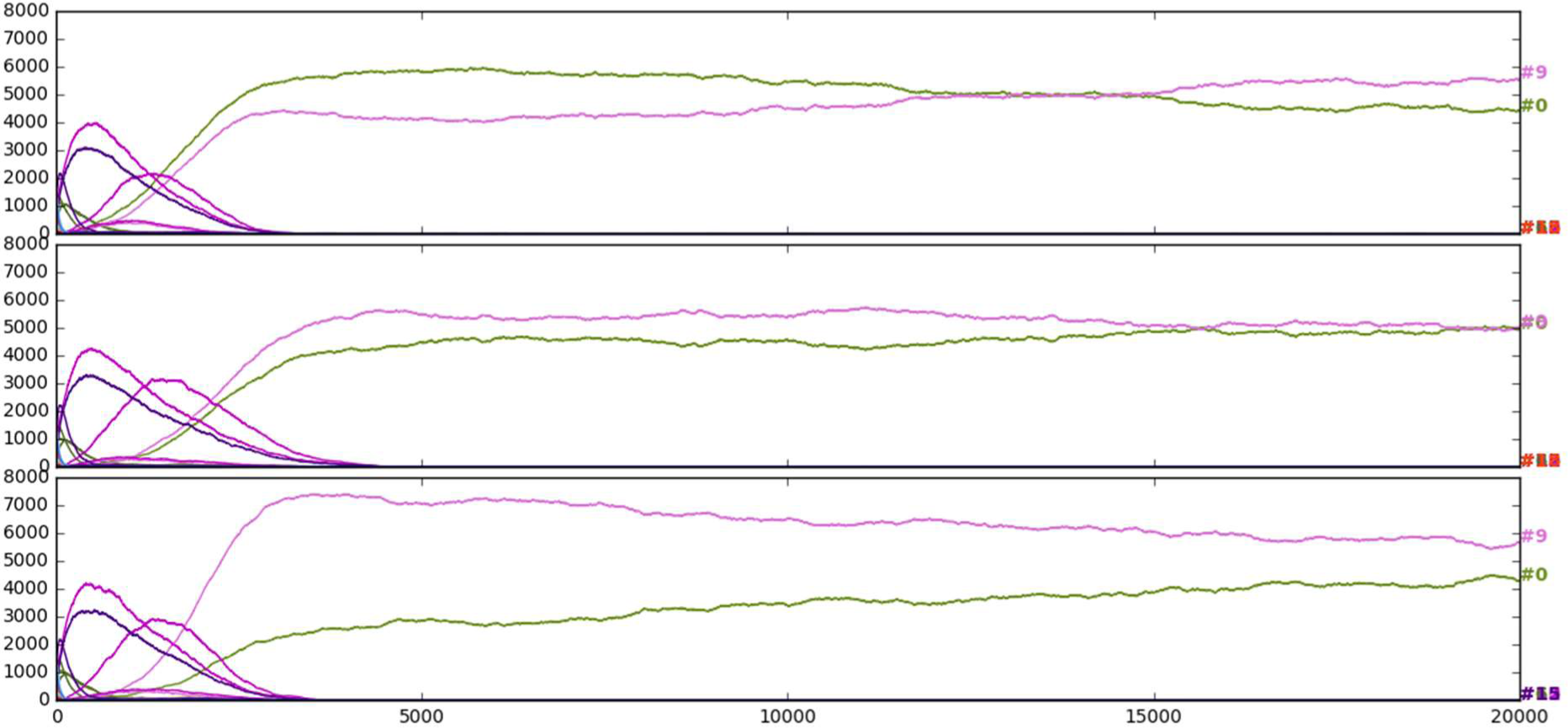
Timeline of three independent runs from region c11, code=387; neighbourhood size 24.

**Figure 10.**
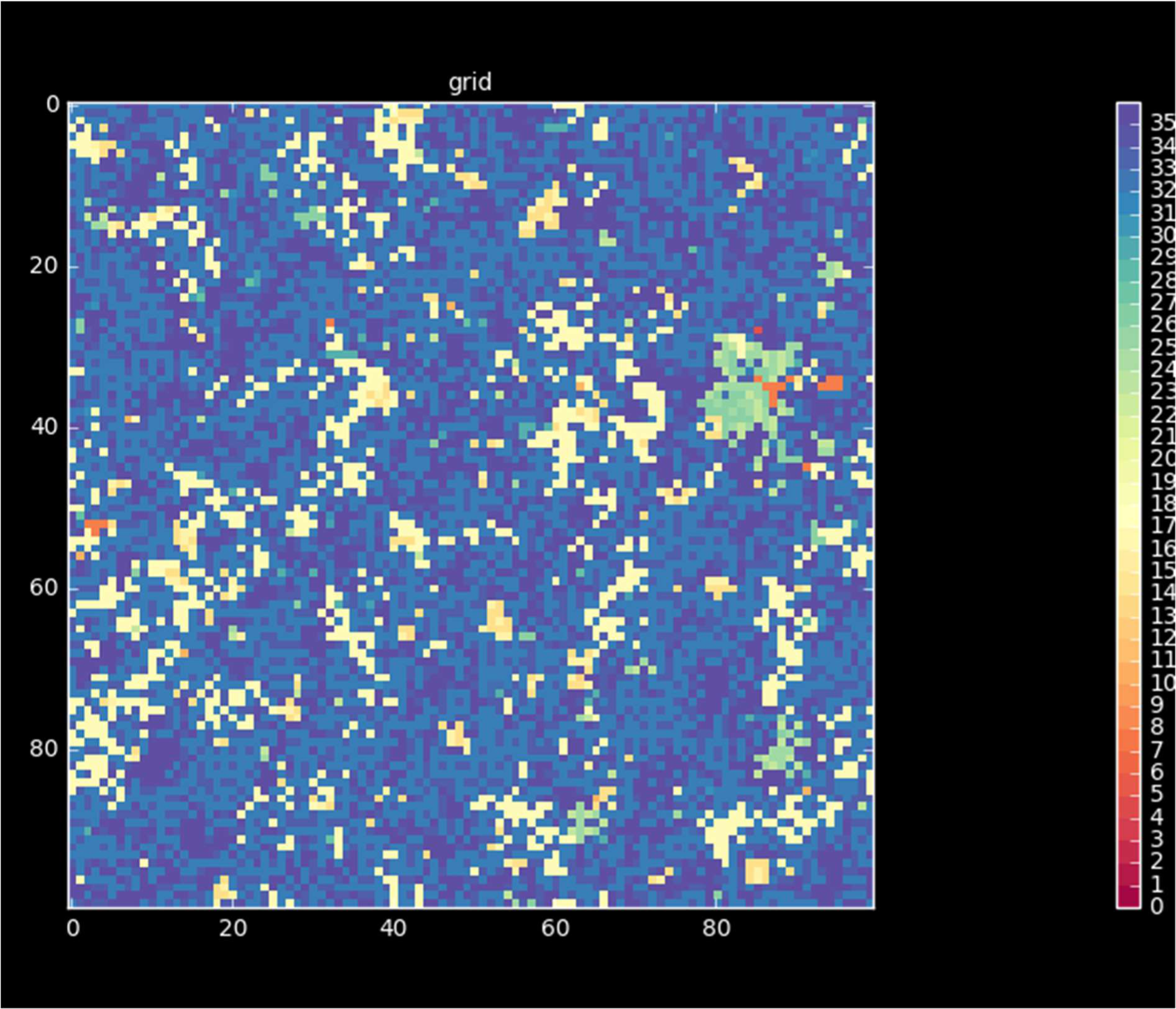
Final grid of the first independent run from region c11, code=8249; neighbourhood size 8.

**Figure 11.**
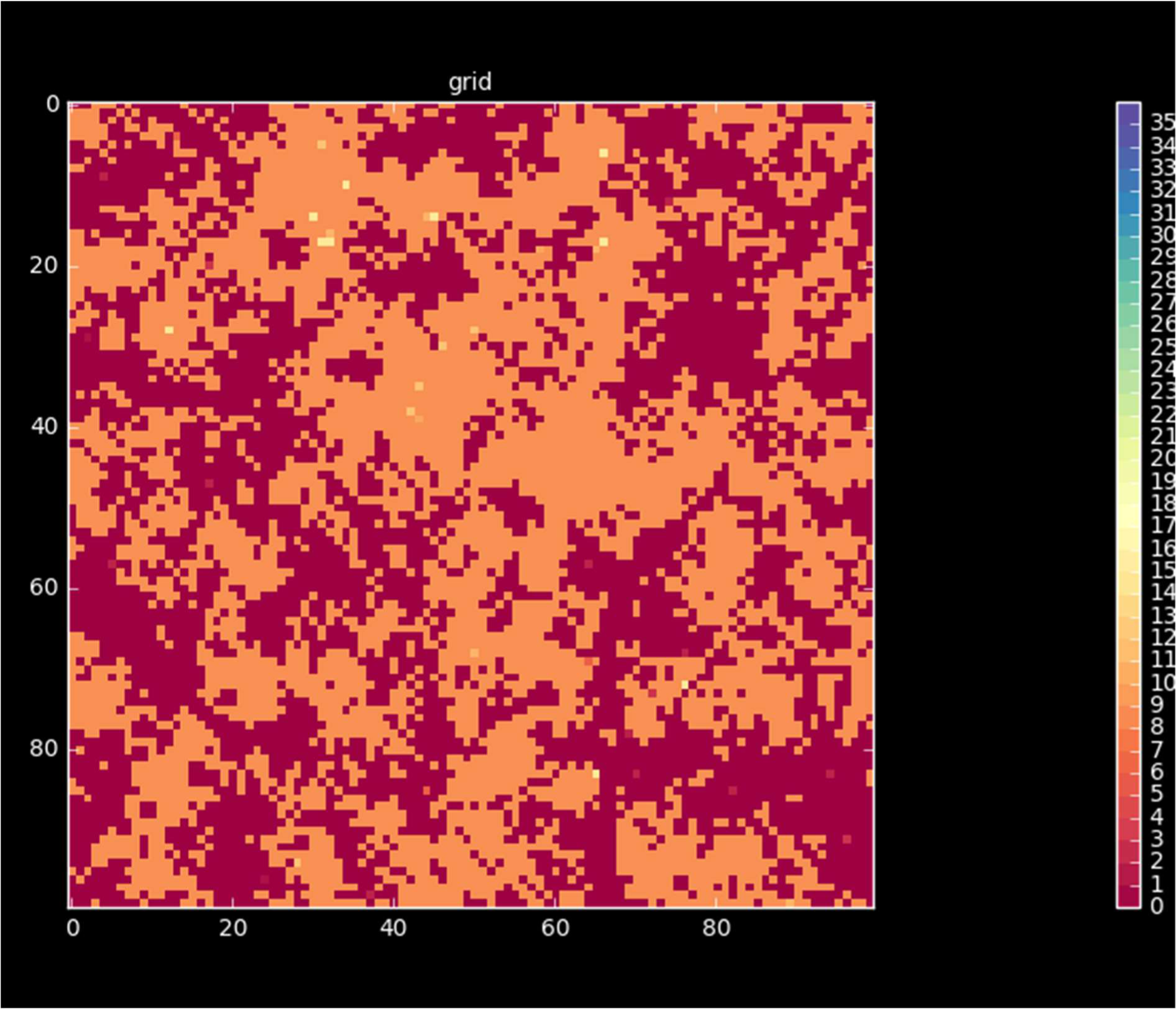
Final grid of the first independent run from region c11, code=387; neighbourhood size 8.

**Figure 12.**
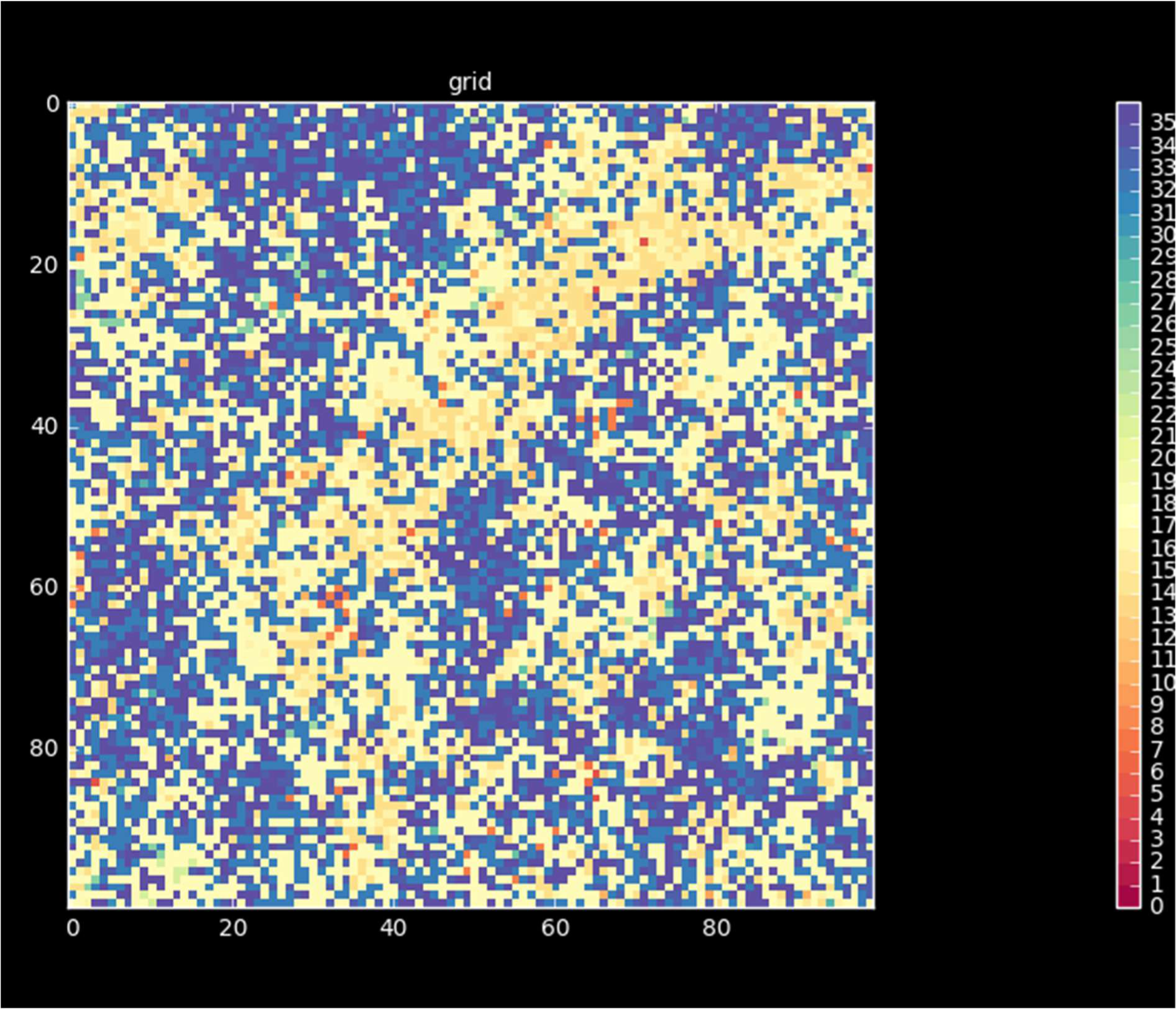
Final grid of the first independent run from region c11, code=8249; neighbourhood size 24.

**Figure 13.**
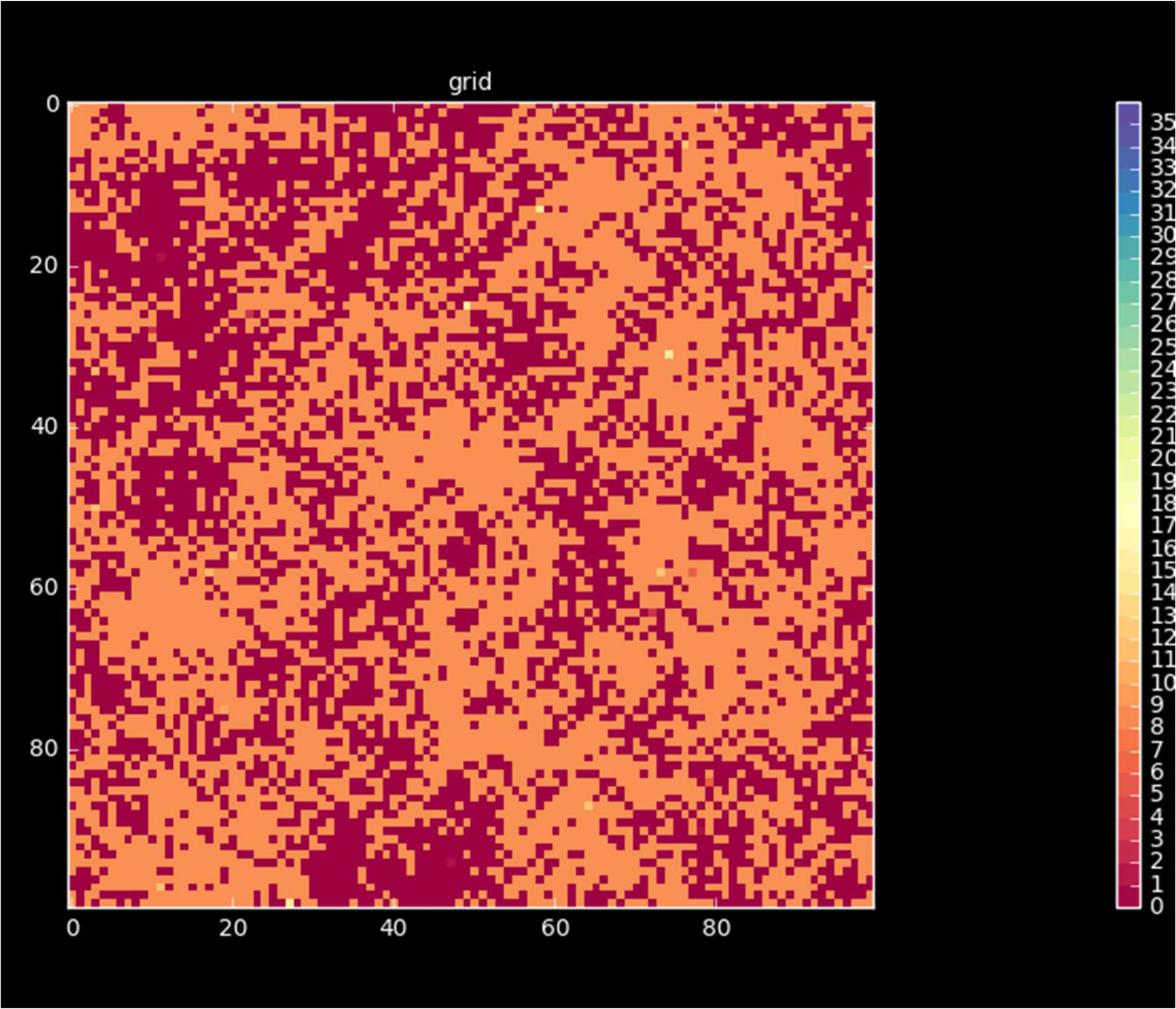
Final grid of the first independent run from region c11, code=387; neighbourhood size 24.

## 4. Discussion

Here we investigated the evolution of honest and cheating strategies in a spatially explicit game of aggressive communication. No honest communication evolved in spatial setting, regardless of the neighbourhood size. Mixed cheating evolved with low neighbourhood size. The result is counterintuitive and requires further study. Spatial correlations help the evolution of cooperation and honest communication is a form of a cooperation. Somewhat similar results were found in the spatially explicit snowdrift game. Hauert and Doebeli [12] found that under some update rules the level of cooperation is smaller in the spatially explicit game than in the well-mixed populations. They concluded that “spatial structure often inhibits the evolution of cooperation” (title). It turned out however that this counterintuitive result is due to the limited neighbourhood size and not due to spatial correlations. A spatial game introduces both spatial correlations and limited range of interactions (which is usually modelled as the neighbourhood size) by definition. Unless the effects can be separated by design it is not correct to single out any of these factors as the source of the effect. The design used by Hauert and Doebeli [12] was not suitable to tell which one of these factors were responsible for the lower level of cooperation. Szamado et al [14] was able to show in a controlled setup that spatial correlations actually do help cooperation in both the PD and SD game.

Changing the neighbourhood size (*n*) also shows a counterintuitive trend. Increasing neighbourhood size should approximate the well mixed results better. One potential problem is that the honest (*S_A_*) global strategy is a mix of weak and strong individuals and in an endogenous model studied here the ratio of weak to strong individuals (*q*) can change as a function of fitness. Since neighbourhood size was limited therefor the number of potential *q* values were also limited depending on n. For example, with *n*=4 the proportion of weak individuals could only take one of the five values: 0, 0.25, 0.5, 0.75, 1. If the equilibrium ratio of weak individuals (*q*\*) is different from any of these values then the honest global strategy will not be an ESS since either weak or strong individuals will disappear from the population (which depends on the fitness advantage of these strategies out of equilibrium). There are two controls for this idea: (i) investigate exogenous populations with fixed ratio of weak to strong individuals, (ii) increase the neighbourhood size even further. Sufficiently large increase should result in a good enough approximation of *q*\*.

## Author contribution

SS conceived the idea, A.S. wrote the IBM, S.S. analysed the results and wrote the paper.

## Founding

S.S. was supported by National Research, Development and Innovation Office (NKFIH) OTKA grant K 108974 and by the European Research Council (ERC) under the European Union’s Horizon 2020 research and innovation programme (grant agreement number 648693). A.S. acknowledges support from GINOP 2.3.2-15-2016-00057 (Az evolúció fényben: elvek és megoldások).

